# InhA inhibitors have activity against non-replicating *Mycobacterium tuberculosis*

**DOI:** 10.1101/2020.08.19.257782

**Authors:** Lindsay Flint, Aaron Korkegian, Tanya Parish

**Affiliations:** Infectious Disease Research Institute, 1616 Eastlake Ave E, Seattle, 98102, USA; Center for Global Infectious Disease Research, Seattle Children’s Research Institute, 307 Westlake Ave N, Seattle, 98109, USA

**Author notes:** Immunotherapy Integration Hub, Seattle Children’s Research Institute, 1100 Olive Way, Seattle, WA 98101, USA. Arzeda 3421 Thorndyke Ave W, Seattle, WA 98119, USA.

**Keywords:** anti-bacterial, bactericidal activity, non-replicating bacteria, anti-tubercular, nutrient starvation

## Abstract

We previously identified a diazaborine series with potential for development as a new tuberculosis drug. This series has activity *in vitro* and *in vivo* and targets cell wall biosynthesis via inhibition of InhA. We tested the ability of two molecules of the diazaborine series to kill non-replicating *Mycobacterium tuberculosis* in the nutrient starvation model; both molecules were bactericidal, reducing viability by >3 logs in 21 days. Activity was not inoculum-dependent and showed similar kill rates to other InhA inhibitors (isoniazid and NITD-916). We conclude that inhibition of InhA is bactericidal against nutrient-starved non-replicating *M. tuberculosis*.

## Manuscript

*Mycobacterium tuberculosis*, the causative agent of tuberculosis, remains a global health scourge with >1 million deaths annually (1, 2). Drug treatment is lengthy, in part because anti-tubercular agents are more effective against rapidly growing bacteria, but much less effective against slowly replicating or non-replicating bacteria. There is a need for new agents that target non-replicating persisters (3). We have previously identified a diazaborine series with potential for development as a new tuberculosis drug (4, 5). This series has activity *in vitro* and *in vivo* and targets cell wall biosynthesis via inhibition of InhA (4). Diazaborine activity does not require a cofactor or require activation by bacterial enzymes (unlike the frontline drug isoniazid which also targets InhA) and so the series has improved properties and a lower frequency of resistance than isoniazid.

We had previously demonstrated that two representative molecules from the series (AN12855 and AN12541) had bactericidal activity against replicating *M. tuberculosis* (4). It has long been suggested that drugs that target the cell wall would not be active against non-dividing bacteria and there is some evidence that the efficacy of isoniazid is reduced under these conditions (6). However, we had previously noted that isoniazid was able to kill non-replicating *M. tuberculosis* in the nutrient starvation model with >3 logs kill in 21 days even at concentrations close to the minimum inhibitory concentration (MIC) (7). We wanted to investigate whether the diazaborines could also kill non-replicating bacteria, which could be a good indicator of their ability to shorten treatment in a novel drug regimen (3).

We tested the ability of two molecules of the diazaborine series in the nutrient starvation model (6). *M. tuberculosis* H37Rv (London Pride: ATCC 25618) (8) was grown to log phase in Middlebrook 7H9 medium containing 10% v/v oleic acid, albumen, dextrose, catalase (OADC) supplement and 0.05 % w/v Tween 80. Bacteria were harvested and resuspended in phosphate-buffered saline (PBS) pH 7.4 plus 0.05 % w/v Tyloxapol for 2 weeks to generate nutrient-starved, non-replicating bacteria (9). Compounds were added (final DMSO concentration of 2%) and CFUs were determined over 21 days by serial dilution and culture on Middlebrook 7H10 agar plates supplemented with 10% v/v OADC for 3-4 weeks. The experiment was carried out twice (independent cultures on different dates). AN12855 demonstrated time-dependent activity against non-replicating bacteria (Fig 1A and 1C) with ∼3 log_10_ kill after 14 days at concentrations equivalent to the IC_90_ (0.090 ± 0.050 µM, n=10 from (4)). AN12541 was similarly active against non-replicating bacteria (Fig 1B and 1D) but with a slightly slower kill rate, reaching ∼2 log kill after 14 days (IC_90_ is 0.11 ± 0.21 µM, n=2 from (4)). There was no outgrowth of resistant mutants, which is sometimes seen with isoniazid after 7 days (7).

**Figure 1.**
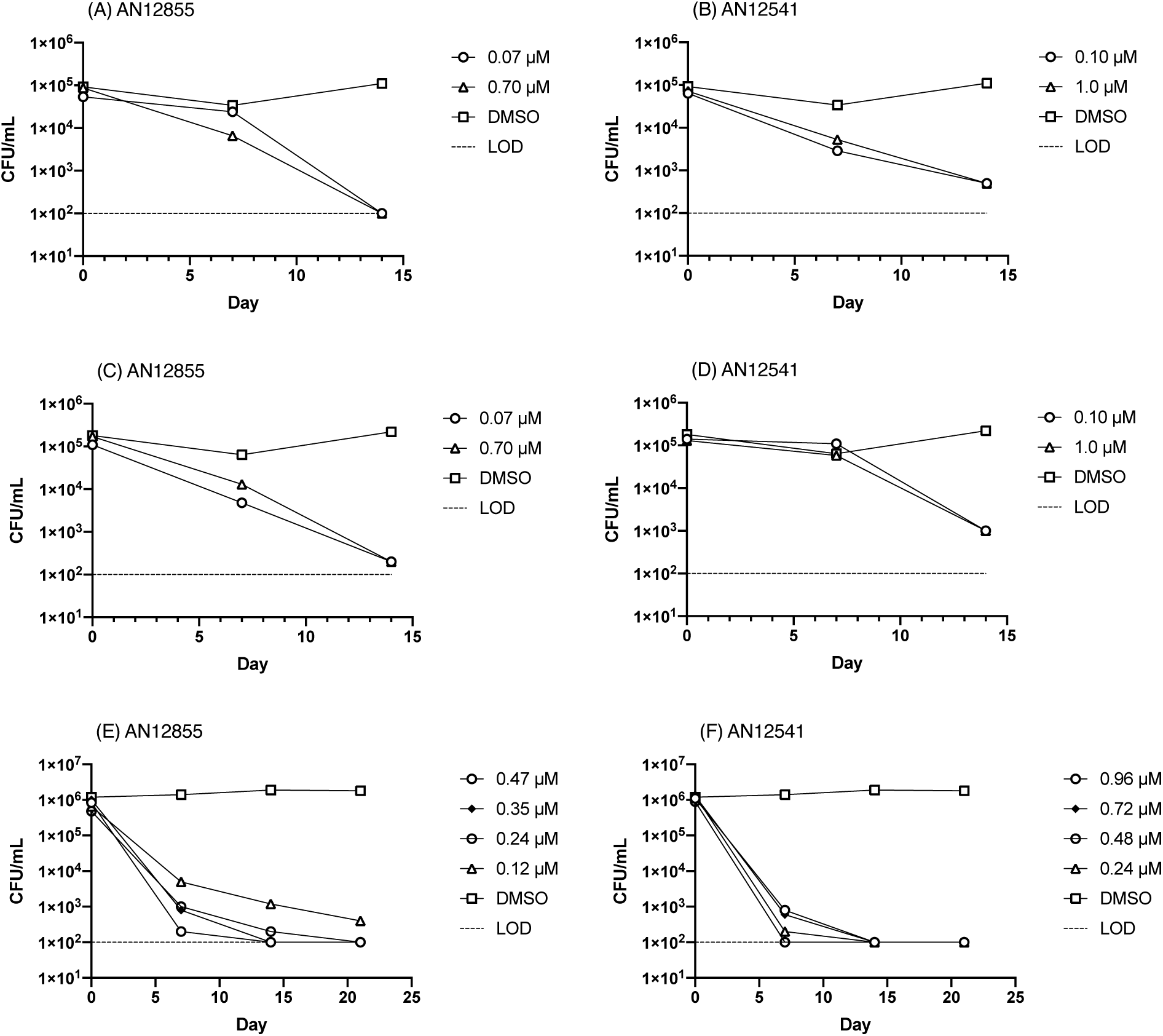
Activity of diazaborines against non-replicating *M. tuberculosis*. Kill kinetics of (A), (C) and (E) AN12855 and (B), (D) and (F) AN12541 against nutrient-starved, non-replicating bacteria. Limit of detection is marked by a dashed line.

Our initial kill kinetics were run with a starting inoculum of 10^5^ CFU/mL to ensure that we did not have any spontaneous resistant clones (which could mask bacterial kill during outgrowth) and over 14 days. We repeated the experiment at a higher inoculum of 10^6^ CFU/mL to determine if there was any inoculum-dependent effect, which has been noted for other anti-tubercular agents with activity against non-replicating *M. tuberculosis* (7). We also used an expanded range of concentrations and an extended period of time (21 days). For AN12855 we saw a similar kill profile over 14 days to our original experiment. Although there was a higher kill rate in the first 7 days and a slower kill rate over the second 7 days, the overall kill rate was similar at 14 days. For AN12541 we saw an accelerated kill rate using the higher inoculum, which was not expected. The difference in the initial kill rate is likely dependent on the physiological state of the bacteria during starvation which may be subtly different between cultures at different densities. Both compounds resulted in >3 logs reduction in viable bacteria over 21 days confirming bactericidal activity (Fig 1E and 1F). There was no outgrowth of resistant mutants at this inoculum.

Isoniazid is inactive against non-replicating *M. tuberculosis* under oxygen limitation and in multi-stress models (10, 11), although we previously demonstrated it had good activity against nutrient-starved bacteria (7). We tested isoniazid and another direct InhA inhibitor, NITD-916 (12), as a comparator for the diazaborine series (Fig 2). We ran two independent experiments (independent cultures on different dates) using the higher inoculum of 10^6^ CFU/mL. Again, we noted that isoniazid had good activity against non-replicating bacteria, as did NITD-916. Both of these showed rapid kill rate with >3 logs kill over 21 days. Kill kinetics were similar between isoniazid and NITD-916. We did not see any outgrowth of resistant mutants for either compound. Thus, we conclude that inhibition of InhA is bactericidal against nutrient-starved non-replicating *M*.*tuberculosis*, and that rapid kill can be effected by InhA inhibitors regardless of the their binding mechanism.

**Figure 2.**
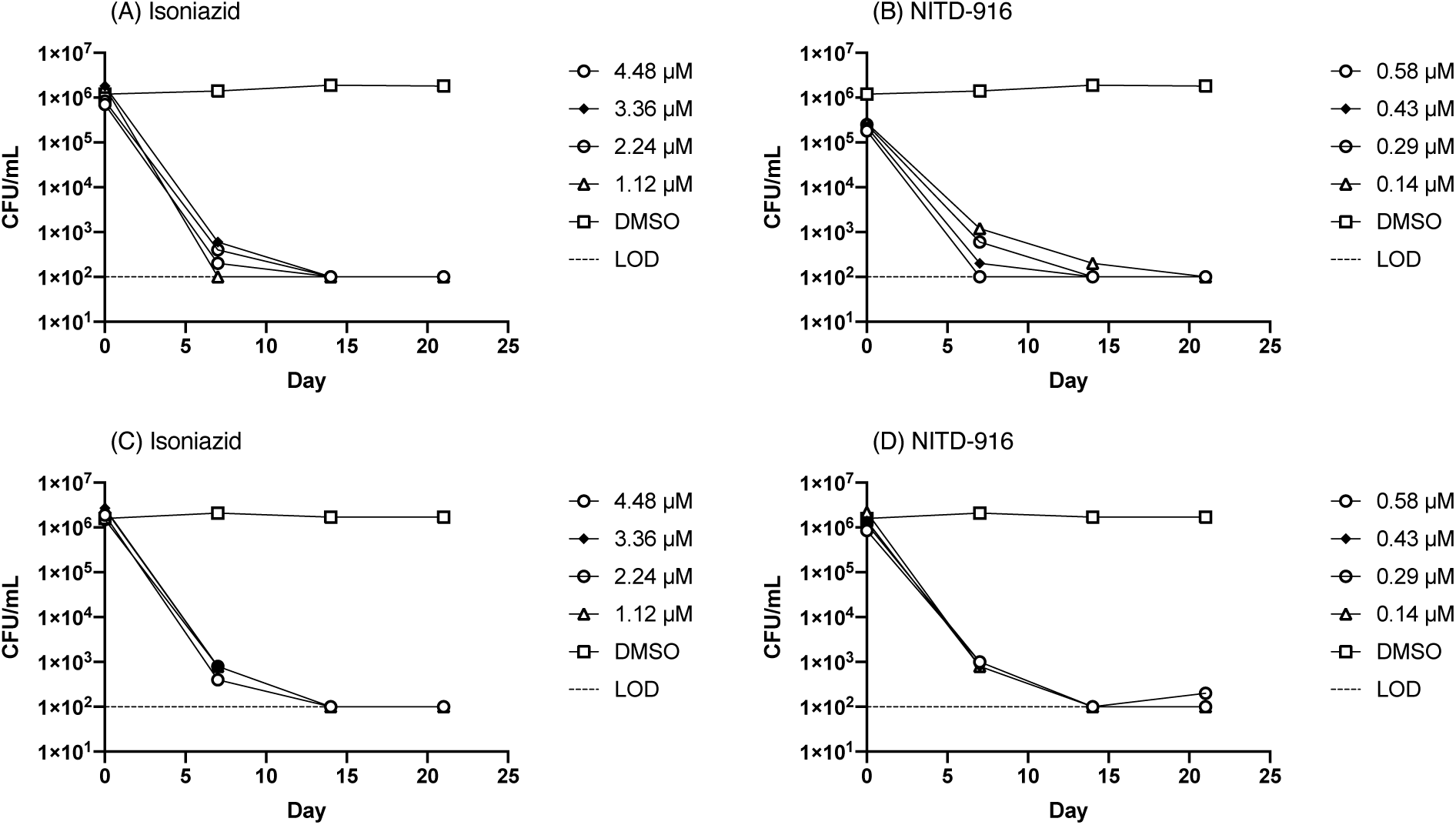
Activity of InhA inhibitors against non-replicating *M. tuberculosis*. Kill kinetics of (A) and (C) isoniazid and (B) and (D) NITD-916 against nutrient-starved, non-replicating bacteria. Limit of detection is marked by a dashed line.

Our work indicates that InhA is a viable target in non-replicating bacteria, at least those generated by starvation. Our data for isoniazid are consistent with those from Betts et al. (6), who demonstrated that isoniazid was able to reduce CFUs in starved bacteria by >1.5 log in 7 days at 10 µg/mL. They did not see activity at 1 µg/mL; however, their starting inoculum was >10^7^ CFU/mL and at that high concentration it is likely that they would have outgrowth of resistant mutants, or that the longer period of starvation used in that study (6 weeks) could be a factor in generating isoniazid resistance.

Taken together with our previous data demonstrating activity against replicating and intracellular bacteria, as well as *in vivo* activity in mouse models of infection (4, 5), these data support the validity of both the target InhA and the diazaborine series for further exploration.

## Authors and contributors

LF and AK conducted the experimental work. LF, AK and TP conceptualized the work and analyzed data. TP wrote the manuscript. LF, AK and TP reviewed and edited the manuscript.

## Acknowledgements

We thank Matt McNeil and Dickon Alley for useful discussion.

